# Causal Dynamics of Social Gaze in Primate Prefrontal-Amygdala Networks Revealed by Dynamic Bayesian Modeling

**DOI:** 10.1101/2025.08.08.669405

**Authors:** Feng Xing, Siqi Fan, Olga Dal Monte, Monika P. Jadi, Steve W.C Chang, Anirvan S. Nandy

## Abstract

Social gaze is a fundamental channel of primate communication, shaping dynamic interactions and fostering mutual understanding. While prior studies have mapped the behavioral correlates of social gaze across the prefrontal-amygdala circuits, the causal architecture of these interactions remains poorly understood. Here, we introduce a novel algorithm to integrate independently recorded sessions into “super-sessions”, validated using ground-truth synthetic data, enabling the reconstruction of simultaneous multi-area recordings aligned to matched gaze sequences. Applying Dynamic Bayesian Network analysis to these super-sessions, we uncover temporally structured, behavior-dependent causal interactions among the amygdala, orbitofrontal cortex, dorsomedial prefrontal cortex (dmPFC), and anterior cingulate cortex. When macaques were the targets of social gaze, the brain-behavior network exhibited positive temporal modulations, with the dmPFC emerging as the dominant source and the amygdala as a primary recipient of influence. When macaques directed their gaze toward their partners, the dmPFC and amygdala retained their respective roles. Prefrontal regions positively modulated one another, while the amygdala acted solely as a downstream target receiving exclusively negatively modulated prefrontal inputs. These findings reveal previously unknown directional interactions in the primate social brain and highlight distinct causal architectures underlying the bidirectional dynamics of social attention.

## Introduction

Gaze behavior plays a crucial role in human cognition, perception, and social interactions (Birmingham et al., 2008; Emery, 2000; Shimojo et al., 2003). Eye-tracking technology enables the investigation of how individuals allocate their visual attention during tasks, providing valuable information about how the brain processes and prioritizes information (Rayner et al., 2009). Understanding how individuals direct their visual attention provides insights into underlying cognitive processes, decision-making mechanisms, and social dynamics (Carrasco, 2011; Corbetta & Shulman, 2002; Land & Tatler, 2009; Posner, 2012). By analyzing eye movements, researchers can infer attentional focus, cognitive workload, and emotional states, making gaze behavior an essential tool for studying both typical and atypical cognitive functions (Emery, 2000; Reingold, 2014). Specifically, gaze behavior is a critical component of social cognition, influencing how people interpret and respond to social cues in communication, joint attention, and empathy (Argyle et al., 1994; de, 2016; Emery, 2000; Frischen et al., 2007; Mundy & Newell, 2007). Beyond fundamental research, gaze behavior analysis has practical applications in clinical domains. Studying gaze patterns has been instrumental in diagnosing and monitoring neurological and psychiatric disorders such as autism spectrum disorder, schizophrenia, and Alzheimer’s disease (Pelphrey et al., 2002; Tseng et al., 2013). Given its broad interdisciplinary significance, studying gaze behavior provides a powerful window into fundamental cognitive mechanisms and offers practical applications across multiple fields.

Nonhuman primates (NHPs) provide a unique advantage in studying gaze behaviors due to their close evolutionary relationship to humans and the similarity of their visual and social processing systems, enabling insights into neural mechanisms underlying complex social cognition (Emery, 2000; Shepherd, 2010). Additionally, experimental approaches such as invasive neural recording and causal manipulations are feasible in NHPs, allowing researchers to directly link neural activity to gaze behaviors in ways not possible in human studies (Fan et al., 2024; Mosher et al, 2014). As technological advancements improve the precision and accessibility of eye-tracking systems, the scope of gaze research continues to expand. Unlike traditional paradigms that primarily relied on static images or prerecorded videos, recent studies have begun to examine live, reciprocal social gaze behavior between pairs of macaques (Dal Monte et al., 2016), allowing for the analysis of interactive and dynamic social exchanges that more accurately reflect naturalistic social cognition.

Investigating the underlying neural mechanisms governing live social gaze behaviors requires examining the key brain regions involved and their intricate interconnections. Notably, regions such as the basolateral amygdala (BLA), gyrus of the anterior cingulate cortex (ACCg), orbital prefrontal cortex (OFC), and dorsal medial prefrontal cortex (dmPFC) are critical components of the primate social interaction network, exhibiting selective neural activation when monkeys observe social interactions among conspecifics (Cléry et al., 2021; Freiwald, 2020; Sliwa & Freiwald, 2017) and playing essential roles in processing social stimuli and guiding appropriate social behaviors across species (Dal Monte et al., 2020; Fan et al., 2024; Gangopadhyay et al., 2021; Magrou et al., 2024; Simon Iv & Rich, 2024; Zeisler et al., 2024). The functional significance of these regions is deeply embedded in their interconnections, supporting the importance of obtaining a network-level understanding of how these brain areas coordinate to guide social gaze interactions. Indeed, anatomical tracing studies have demonstrated robust bidirectional projections between the BLA and prefrontal regions, facilitating the integration of emotional and social information necessary for adaptive behavior (Ghashghaei et al., 2007; Kennedy & Adolphs, 2013). The OFC and dmPFC are reciprocally connected both with each other and with the ACCg, supporting flexible decision making, social evaluation, and monitoring of outcomes (Carmichael & Price, 1995; Joyce et al., 2025; Petrides & Pandya, 1999). These extensive anatomical connections enable coordinated processing of affective and social cues, underscoring the importance of distributed circuits in primate social behavior.

A recent study has found that large proportions of neurons in the prefrontal cortical areas and the amygdala display diverse temporal response patterns for monitoring either self- or other-directed gaze and for selectively signaling mutual gaze events, highlighting the extensive presence of interactive social gaze coding throughout these networks (Dal Monte et al., 2022). However, this study only examined each brain region independently, without addressing how these regions interact or causally coordinate their activity to guide live social gaze interaction. Understanding causality in neuroscience is essential for deciphering the mechanisms underlying brain function and behavior. Traditional correlation-based analyses, while useful, often fail to capture true causal relationships between neural activity and cognitive or behavioral outcomes (Pearl, 2009; Reid et al., 2019). Dynamic Bayesian Networks (DBNs) offer a powerful probabilistic framework for modeling temporal dependencies in multivariate neural systems. DBNs explicitly represent how a time-series evolves, making them particularly suited for studying causal interactions in neural data (Murphy, 2002). By leveraging their ability to infer causal relationships from sequential data, DBNs have been successfully applied in neuroscience to model brain connectivity and predict neural responses to stimuli (Das et al, 2024).

Here, we developed a novel algorithm to merge separately recorded sessions into unified “super-sessions”. This approach enables the simulation of simultaneous recordings from multiple brain regions precisely aligned to matched behavioral sequences. Using an analysis approach based on Dynamic Bayesian Networks, we identified temporally specific causal interactions between social gaze behavior and multiple brain areas in the prefrontal-amygdala networks, including the BLA, ACCg, OFC, and dmPFC. We find that receiving directed attention from a partner consistently increased network dependencies, particularly with dmPFC as the dominant source and the BLA as the dominant target. In contrast, when gazes were directed toward the partner, prefrontal areas exhibited mutual positive modulation, whereas the amygdala served strictly as a downstream node receiving only negative dependency modulations from prefrontal regions. Our approach uncovers previously unrecognized causal dynamics among the social brain network during naturalistic gaze interactions, underscoring the importance of elucidating the causal architecture underlying social cognition.

## Results

### Experimental setup and social gaze categories

We analyzed recently published data (Dal Monte et al., 2022) in which pairs of rhesus macaques (‘Actor’: recorded monkey; ‘Partner’: partner monkey; Fig. 1A) freely interacted with one another using gaze in a face-to-face setting. Eye-tracking was performed simultaneously in each monkey with high temporal and spatial resolution (see Methods). The BLA and one of the three prefrontal areas – ACCg, dmPFC, OFC – were simultaneously recorded in each recording session (Fig. 1B). Spiking activity was recorded from 537 BLA, 236 ACCg, 187 dmPFC, and 241 OFC neurons from two actor monkeys.

**Figure 1.**
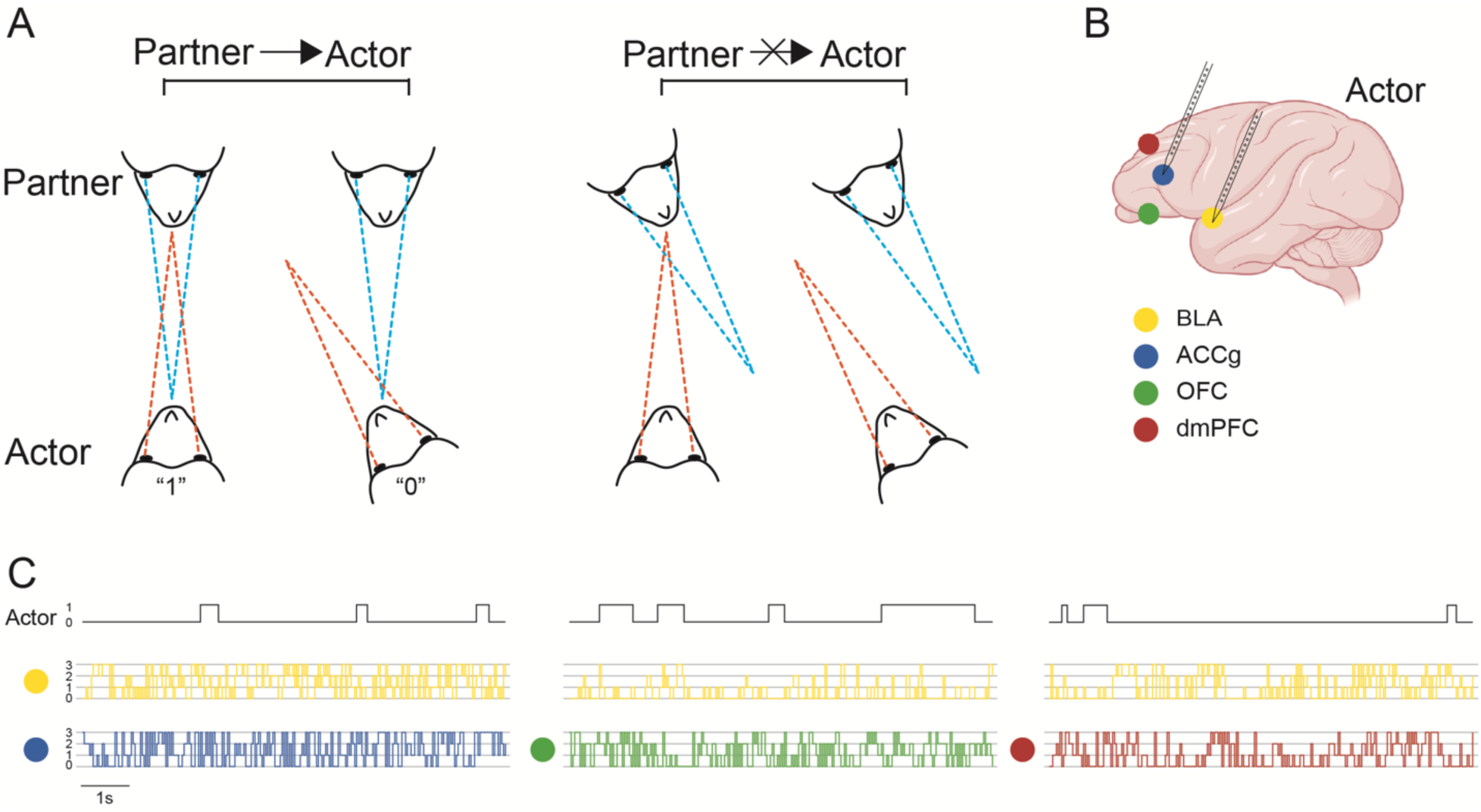
Social gaze interactions and example traces of gaze and neural variables. (**A**) Two monkeys, designated as “actor” and “partner”, engaged in free-viewing face-to-face social gaze interactions. Two conditions are depicted: partner gazing at actor’s face (left), partner looking elsewhere (right). Actor’s gaze variable was coded as “1” (actor gazing at partner’s face) or “0” (actor not gazing at partner’s face). (**B**) Actor’s recording sites: basolateral amygdala (BLA), anterior cingulate cortex gyrus (ACCg), orbitofrontal cortex (OFC), and dorsomedial frontal cortex (dmPFC). In every recording session, the BLA and one other area among ACCg, OFC, and dmPFC were simultaneously recorded. (**C**) Example traces of actor’s gaze and neural variables. Actor’s neural variables are categorized into four quantile levels (0,1,2,3). Colors are same as in **B**.

Based on the previously established significance of gaze toward the face as a strong social stimulus in both humans and non-human primates (Dal Monte et al., 2016; Gothard et al., 2007; Itier & Batty, 2009; Kano et al., 2018), we considered the face region of a conspecific as the main region of interest (ROI) in our study. Specifically, we were first interested in understanding how the brain-behavior network in the actor animal was modulated by social gaze from the partner animal. Experimental sessions were therefore partitioned into specific behavioral epochs based on whether the partner gazed at the actor’s face (Partner → Actor) or looked elsewhere (Partner ↛ Actor) (Fig. 1A). Within each of these epochs, the actor’s gaze variable was set to “1” when it gazed at the partner’s face; otherwise, it was set to “0”.

Example traces of the actor’s gaze variable and neural variables are shown in Fig. 1C. To normalize the varying degrees of single-unit yield across recording days, we categorized the aggregated spiking activity in each brain region using quantiles into four levels (0,1,2,3) for all subsequent analyses (see Methods).

### An algorithm for discovering causal dependencies in “super-sessions”

To “super-session”, or simulate simultaneous recordings, we assume that the underlying neural dynamics within the brain network of interest remain consistent across identical patterns of the pertinent behavior, specifically gaze, regardless of whether neural activity is recorded. Based on this premise, we merged neural data across recording sessions by aligning them to matched gaze patterns and proceeded to identify dependencies across time lags that are conditioned on the entire brain network of interest. The key challenge in discovering conditional dependencies in multivariate data that is partially observed across sessions is the detection of spurious dependencies. To mitigate this, we used an analysis approach that fits probabilistic graphical models to estimate a confidence measure of conditional dependencies at multiple time lags (Das et al., 2024).

We first demonstrate our approach in ground truth synthetic data created from three-component linear (Fig. 2A) and nonlinear (Fig. 2B) systems. In both systems, X_1_ is a random binary time series, X_2_ is a function of X_1_, and X_3_ is a function of both X_1_ and X_2_. We then generated paired data of {X_1_, X_2_} and {X_1_, X_3_} by simulating the underlying systems (Fig. 2C). By aligning data to X_1_ as the anchor variable (shown in the same outlined color in Fig. 2C), we reconstructed full datasets of {X_1_, X_2_, X_3_}, separately for the linear and nonlinear systems. We refer to these reconstructed datasets as “super-sessions”. Our analysis pipeline (see Methods) yielded a weighted dependency matrix, which reveals the dependence structure between network nodes at different time lags. The dependency matrix, depicted in Fig. 2D for the linear system, accurately captured the relationships between variables at the correct temporal lags (compare Fig. 2D and the linear system in Fig. 2A).

**Figure 2.**
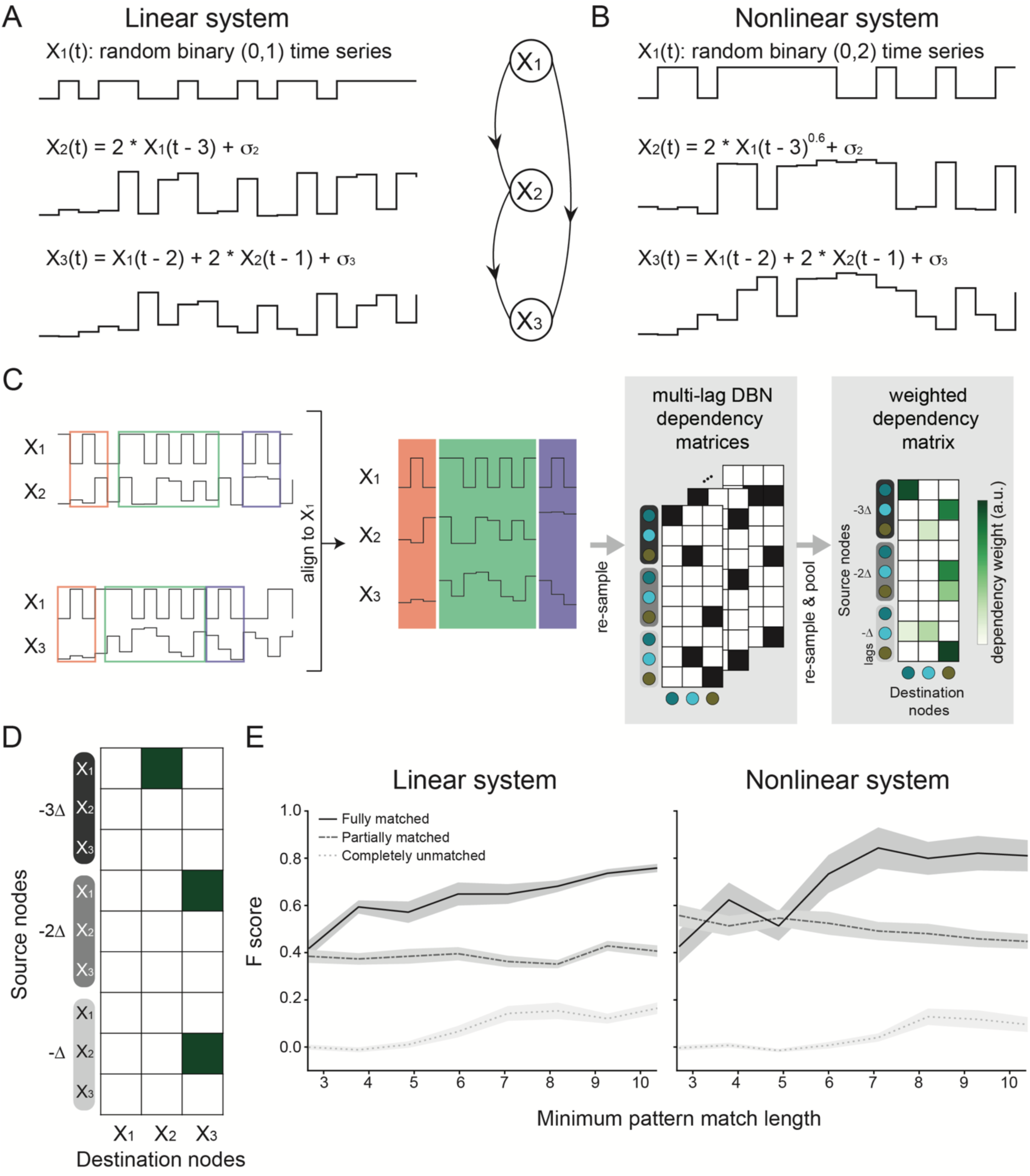
Validation of “super-sessioning” method on ground truth simulations of linear and non-linear systems. (**A**) A three-component linear system, consisting of variables X_1_, X_2_, and X_3_. X_1_ is a random binary time series and serves as the anchor variable. X_2_ is a function of X_1,_ while X_3_ is a function of X_1_ and X_2_. The formulas show the exact relationships. Example traces are shown. σ_2_ and σ_3_ are Gaussian noise terms. (**B**) A three-component non-linear system, similar in structure to A, except X_2_ is a non-linear function of X_1_. (**C**) Pipeline of the merging algorithm and Dynamic Bayesian Network (DBN) based discovery of causal dependencies. Pairs of {X_1_, X_2_} and {X_1_, X_3_} data were aligned to patterns in X_1_. The same color outline refers to matched patterns of X_1_ in partial data. The merged super-session was re-sampled and served as input to the DBNs pipeline (see Methods) to generate a weighed dependency matrix. (**D**) Weighted dependency matrix from the simulated linear system. The green squares indicate dependency from a source node (Y axis) to a destination node (X axis). Δ indicates time lags. (**E**) F-score metric as a function of minimum pattern matching length for the linear and non-linear systems. The solid, dashed, and dotted lines refer to the fully matched, partially matched, and completely unmatched groups respectively. The gray shading represents the standard error of the mean (n = 100).

To test if the merging algorithm was sensitive to the length of the matched patterns, we calculated a measure of similarity (F-score) between the DBN-discovered dependence structure and the ground truth structure for different minimum match lengths (Fig. 2E; see Methods). Crucially, as controls, we calculated the F-scores for partially matched and completely unmatched datasets (see Methods). For both linear and nonlinear systems, the fully matched group outperformed the partially matched and completely unmatched groups, reflecting that DBN can reliably discover dependencies from super-sessions regardless of linearity. The F-scores for the partially matched group were higher than those for the completely unmatched group, indicating that DBN can discover some dependencies even from partial information. For the linear system, the F score increased monotonically with longer minimum match lengths, indicating that longer matched patterns contribute more robustly to the dependence inference process. For the nonlinear system, the F-score rose with increasing match length but quickly saturated at a high score, suggesting that moderately long patterns are sufficient to capture the underlying dependencies, with limited gains from extending the match further. The F scores of the linear and nonlinear systems were comparable across groups with similar matching conditions, demonstrating the effectiveness of the DBN framework in both contexts. However, the nonlinear system exhibits greater variability under both fully and partially matched conditions, indicating that dependency inference in nonlinear systems is more susceptible to noise and variability in the input data.

In addition to validating the super-session approach with ground truth simulations, we also performed validation tests on the empirical data. Since BLA activity was always recorded daily with one of the prefrontal areas, we swapped BLA activity across recording days by matching the actor’s gaze patterns under various partner gaze conditions (Fig. S1; see Methods). As shown in Fig. S1A, when the partner gazed at the actor, the F-scores of the BLA-swapped group were significantly higher than those of the randomized BLA-swapped control group (Fig. S1A; Mann-Whitney test, p < 10⁻⁴⁰). This result remained consistent when the partner did not gaze at the actor (Fig. S1B; Mann-Whitney test, p < 10⁻³⁸).

Our method thus allowed us to merge separate experimental sessions into super-sessions based on the matched patterns of one common variable – the actor’s gaze – to discover a more comprehensive causal brain-behavior network during social gaze interactions.

### Causal interactions in the actor’s brain-behavior network are upregulated by partner’s social gaze

Using matched patterns of the actor’s gaze variable, we generated super-sessions of actor gaze behaviors and neural recordings from BLA, ACCg, OFC, and dmPFC (Fig. 3A) under different partner gaze conditions (Fig. 1A). Super-sessions were resampled and prepared into data tables (Fig. 3B). We chose four time-bins of 50ms each—current time bin (‘0’) and 3 lagged bins (−Δ, −2Δ, −3Δ)—to cover a timespan of 200ms as the basic unit of analysis. This timespan was chosen to match the canonical timescales of fixation in primates (König & Buffalo, 2016). The matched behavior epoch durations were best described by an exponential decay with a time constant of 300.5 ms (Fig. S2; see Methods). The data tables served as inputs to the DBN analysis framework (Fig. 2B) to yield weighted dependency matrices (Fig. 4A-B). Dependency weights were higher in the Partner → Actor condition (Fig. 4A) compared to the Partner ↛ Actor condition (Fig. 4B), supporting the sensitivity of the actor’s brain-behavior network to partner’s social engagement. To quantify the impact of social gaze from the partner on the actor’s brain-behavior network, we calculated the modulation of the dependency weights (for only those edges that survived a time shuffle control; Methods) to generate a modulation matrix (Fig. 4C). Notably, all the dependency modulations were positive, indicating an overall enhancement of causal interactions in the actor’s brain-behavior network due to social gaze from the partner.

**Figure 3.**
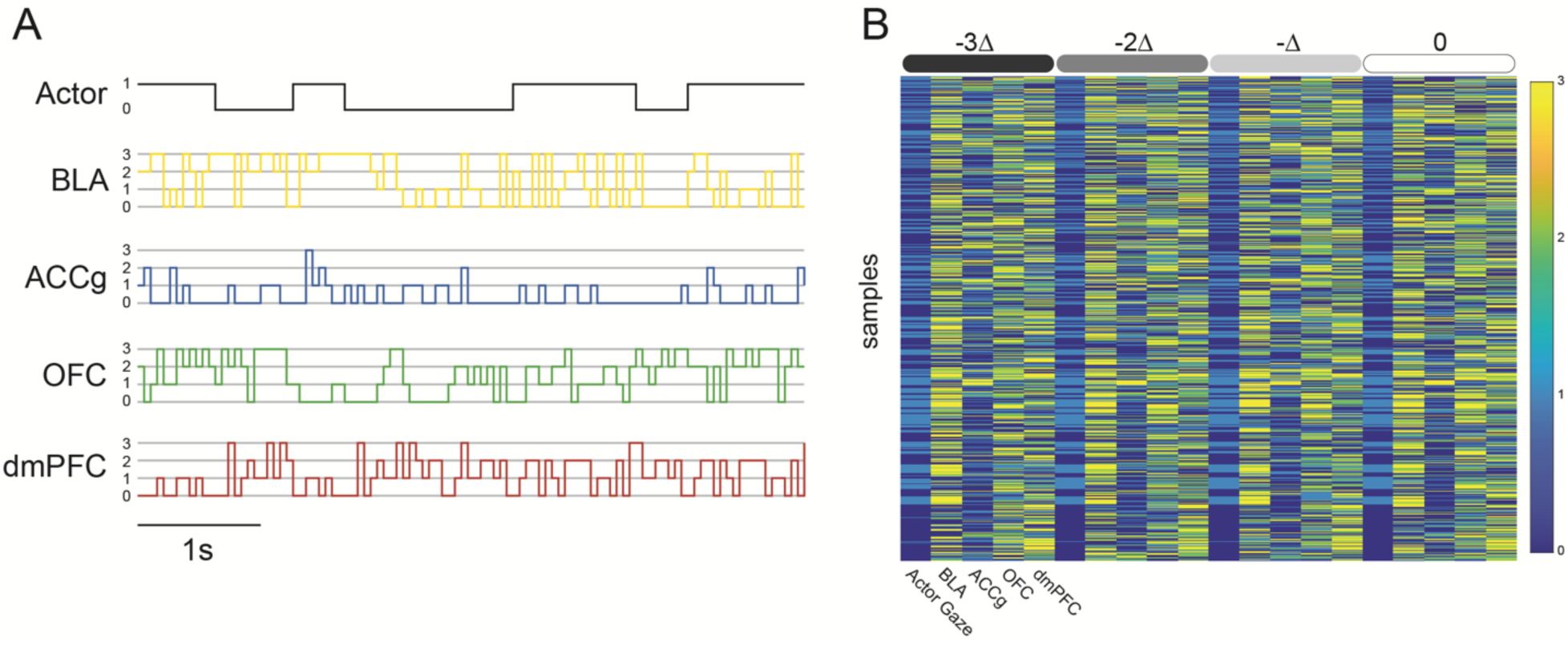
Example of merged super-session data and prepared data table for DBN analysis. (**A**) Example traces of merged super-session based on matched gaze patterns of the actor. (**B**) Prepared data table from merged super-session. The bars on top denote data columns at different time lags. The first column of every time lag represents the actor’s gaze variable and the remaining columns are neural variables. Δ=50ms.

**Figure 4.**
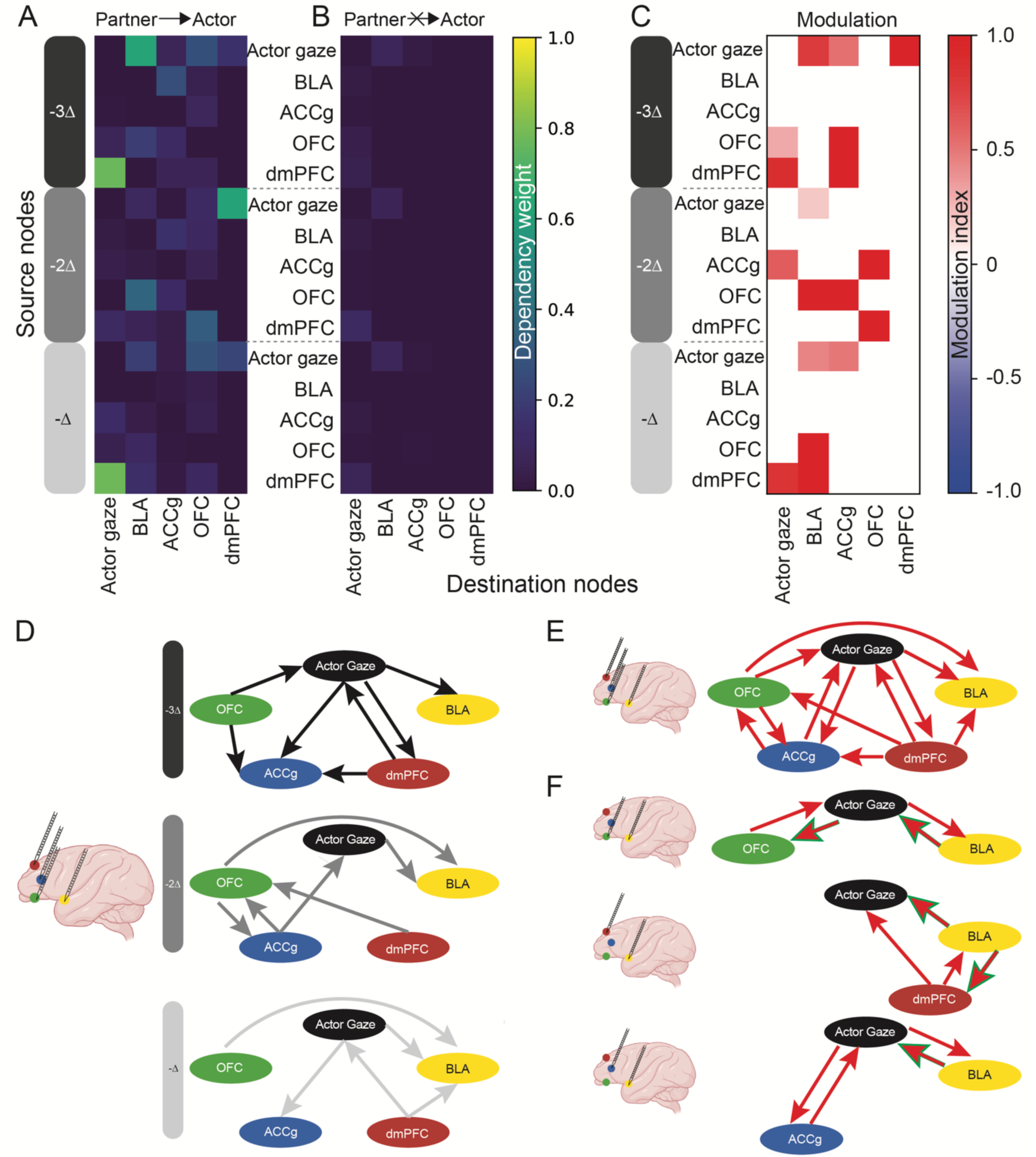
Causal interactions in the actor’s brain-behavior network due to partner’s social gaze. (**A**) Weighted dependency matrix for the Partner → Actor gaze condition. Y axis indicates source nodes and X axis indicated destination nodes. Different colored bars along Y axis denote different time lags. (**B**) Weighted dependency matrices for the Partner ↛ Actor condition. (**C**) Dependency modulation matrix between the two conditions. (**D**) Modulation maps at different time lags. The edges indicate the direction of the dependency modulation. The edges are color-coded for the time lags. (**E**) Merged comprehensive modulation map obtained by merging the delay-resolved maps in D. The edges indicate the direction of the dependency modulation. The edges are color-coded for the sign of modulation (red, positive modulation). (**F**) Partial modulation maps were generated from the dual area recordings without super-sessioning. The green outlined edges indicate newly emerging edges in the partial maps that are not present in the comprehensive map. Δ=50ms.

To further elucidate the nature of the enhancements, we visualized the modulation matrix as modulation maps at different time lags (Fig. 4D). These maps illustrate the modulation of the causal linkages in the network at different time scales. Actor’s gaze consistently had positive dependency modulation directed toward the BLA from the prefrontal regions. Previous studies have revealed the general role of the amygdala in attending to faces and detecting their emotional signals (Phelps & LeDoux, 2005; Santos et al., 2011; Tsao et al., 2003). However, it was unknown how the neural activity in the amygdala was modulated when being attended by others. Our finding here demonstrates this modulatory function of BLA from the perspective of causal networks. Further, the ACCg and dmPFC had bidirectional dependency modulation with Actor’s gaze at earlier time points, which can explain the earlier emergence of selectivity for mutual eye contact in these areas compared to the BLA and OFC (Dal Monte et al., 2022).

To gain an understanding of the overall connectivity modulations of the brain-behavior network, we next examined the structure of a comprehensive modulation map by collapsing across the maps at different time lags (Fig. 4E). We found that the BLA was the recipient of dependency modulation from other prefrontal areas, reflecting that the function of BLA is shaped by neural representations of its interconnected prefrontal areas (Murray & Fellows, 2022). In contrast, the dmPFC was the source of dependency modulation to all the other nodes tested, implying the crucial role of dmPFC in the brain-behavior network, which is in agreement with the dmPFC’s essential role in “Theory of Mind” (Báez-Mendoza & Williams, 2020; Hayashi et al., 2020; Iacoboni et al., 2005; Martin et al., 2017; Martin et al., 2021).

Furthermore, to validate whether the comprehensive modulation map obtained from super-sessioning accurately captured the connectivity of the brain-behavior network, we compared this comprehensive map to those generated from the simultaneous dual area recordings (partial dependency modulation maps; Fig. 4F). Although there were some newly emerged edges (outlined in green in Fig. 4F) in the partial dependency modulation maps, they can be explained by the structure of comprehensive map. For instance, the edge from Actor gaze to OFC in the {Actor gaze, OFC, BLA} map (Fig. 4F, top) can be explained by edges originating from the missing dmPFC node to the Actor gaze and OFC nodes in the comprehensive map. This analysis thus provides further confidence that the comprehensive map from the super-session reveals the dynamics of the actor’s brain-behavior causal network due to social gaze from the partner.

### Prefrontal-amygdala network is differentially modulated by social gaze context

Being gazed at by others versus gazing at others both activate neurons in the prefrontal cortex and the amygdala (Dal Monte et al., 2022). From the perspective of the actor, these two contexts are distinct in terms of the directionality of social gaze and hence of social agency and awareness. We hypothesized that the prefrontal-amygdala network would be differentially modulated by these two critical social contexts. To test this, we investigated network interactions under the following non-reciprocal gaze conditions: directing gaze to the partner (Actor → Partner) and receiving gaze from the partner (Partner → Actor) . The corresponding weighted dependency matrices are shown in Fig. 5A. Notably, the two matrices are substantially different, reflecting different network interaction structures under the two contexts. We then calculated the dependency modulation matrix (for only those edges that survived time-shuffled control, see Methods; Fig. 5B) and plotted the dependency modulation maps at different time lags (Fig. 5C). Notably, dependency modulations were consistently biased in sign across regions: BLA received only negative modulations, whereas ACCg and OFC primarily received positive ones. As observed previously, BLA was exclusively modulated by prefrontal sources, but all incoming modulations were negative when the actor directed their gazes at the partner. In contrast, ACCg and OFC were positively modulated by other prefrontal areas. Specifically, ACCg received positive dependency modulations from OFC and dmPFC at longer lags (−3Δ, −2Δ) when the actor gazed at the partner. Likewise, OFC was positively modulated by ACCg and dmPFC at shorter lags (−2Δ, −Δ), with only a marginally negative modulation from dmPFC at −2Δ. Once again, dmPFC emerged as the sole source of all dependency modulations, reinforcing its upstream function. Together, these results reveal a hierarchical organization of dependency modulations within the prefrontal–amygdala network during social gaze interaction. Prefrontal areas were positively modulated by other prefrontal inputs, with ACCg influenced earlier than OFC. BLA, which only acted as a recipient, received exclusively negative modulations from prefrontal areas, highlighting its unique downstream role in this social neural circuit.

**Figure 5.**
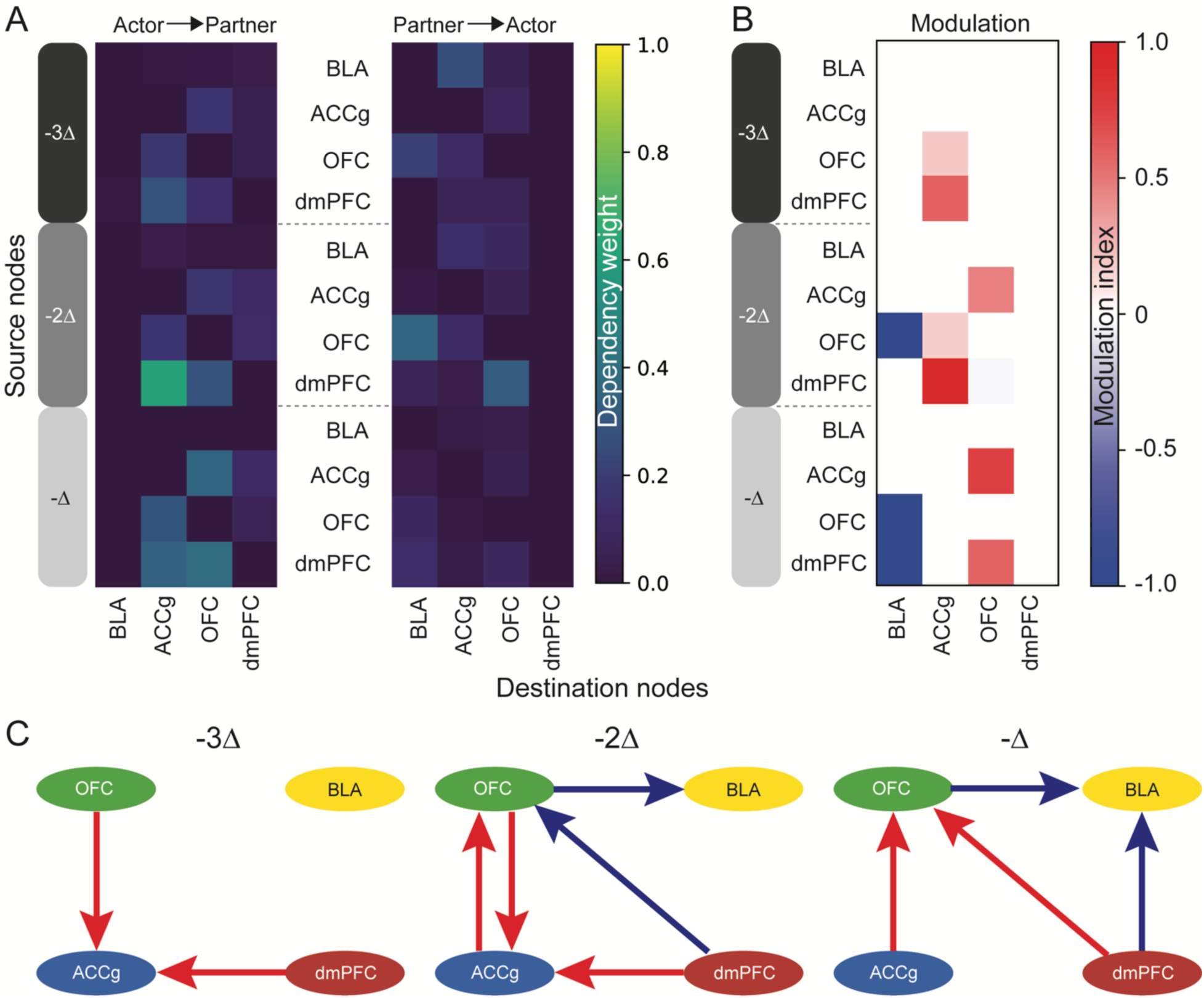
Causal interactions in the prefrontal-amygdala network under different gaze contexts. (**A**) Weighted dependency matrices for the gaze toward partner (Actor → Partner) and gaze from partner (Partner → Actor) conditions. Y axis indicates source nodes and X axis indicated destination nodes. Different colored bars along Y axis denote different time lags. (**B**) Dependency modulation matrix between the two conditions (Actor → Partner versus Partner → Actor) from one area to another. (**C**) Modulation maps at different time points. The edges indicate the direction of the dependency modulation. The edges are color coded as the sign of the modulation (red=positive modulation, blue=negative modulation). Δ=50ms.

## Discussion

The present study demonstrates the effectiveness of a novel merging algorithm in reconstructing brain-behavior networks during unconstrained, naturalistic, behaviors by aligning neural data based on matched patterns of a critical behavioral variable. Through simulations of both linear and nonlinear systems, in which the ground truth is known, we verified that the algorithm successfully merges partial datasets into comprehensive super-sessions, revealing intricate causal dependencies between network variables underlying the modeled dynamical systems. The algorithm’s robustness was reflected in the weighted dependency matrices, which accurately captured both direct and time-lagged relationships, aligning with theoretical expectations (Das et al., 2024). Furthermore, the performance of the merging algorithm, as assessed by the F-score, indicated that even partial matches contribute to network reconstruction, though complete matches provide the highest accuracy. Notably, the nonlinear system exhibited saturation in F-score improvement with increasing match length, suggesting an optimal range for pattern matching before reaching diminishing returns. The comparable F scores observed between linear and nonlinear systems under matched conditions indicate that the DBN framework generalizes effectively across systems with differing underlying dynamics. Nevertheless, the nonlinear system consistently demonstrates higher variability than the linear system across both completely and partially matched conditions, implying that dependency inference in nonlinear dynamics may be more sensitive to noise and less stable under comparable conditions. These findings highlight the utility of the merging algorithm in integrating fragmented data for dynamic network modeling.

Applying this algorithm to brain-behavior data under different gaze conditions allowed us to uncover distinct causal linkages across cortical regions that play a critical role in social cognition. When Partner gazed at Actor, the dependency modulation maps revealed an overall enhancement in brain-behavior interactions, particularly within the BLA, ACCg, dmPFC, and OFC, which are key nodes in the primate social brain that interface with face-selective regions in the earlier visual processing stream (Cao et al., 2025; Kanwisher, 2000; Tsao et al., 2008). The strong modulation observed in BLA aligns with its well-established role in processing social attention and facial expressions (Gothard et al., 2007; Phelps & LeDoux, 2005; Santos et al., 2011; Tsao & Freiwald, 2003). The bidirectional modulation between ACCg and dmPFC at early time points provides further support for previous findings that these regions encode mutual eye contact before amygdala involvement (Dal Monte et al., 2022).

Anatomical studies (Carmichael & Price, 1995; Freese & Amaral, 2005; McDonald, 1998) have shown that the BLA, ACCg, OFC, and dmPFC are extensively interconnected. Functionally, previous work has primarily reported interactions between region pairs. For example, Liu et al. demonstrated that chronic stress alters the excitatory/inhibitory balance in dmPFC neurons projecting to the BLA, which was associated with anxiety-like behaviors (Liu et al., 2023). Additional pairwise interactions have been documented between BLA and OFC (Schoenbaum & Shaham, 2008), ACCg and OFC (Rudebeck & Murray, 2014), and ACCg and dmPFC (Amodio & Frith, 2006). Here our findings reveal for the first time causal dependencies between behavior and this full set of regions (Fig. 4D, 4E), showing how the presence of social gaze—when the Actor is being gazed at— positively modulates interregional interactions, suggesting a dynamic up-regulation of the brain-behavior network in response to social attention cues.

Our results provide new insights into prefrontal-amygdala interactions during different gaze contexts. We find significant differences in network structure when Actor gazed at Partner versus when Actor was gazed at. In particular, prefrontal nodes exhibited asymmetric and temporally organized dependency modulations, revealing functional heterogeneity across regions involved in social cognition and putative differences in computational roles across the social brain. Specifically, ACCg and OFC primarily received positive modulations from other prefrontal areas, consistent with the view that excitatory signaling underpins intra-prefrontal communication during social monitoring (Apps et al., 2016; Lockwood et al., 2016). In contrast, BLA acted exclusively as a recipient and received only negative modulations from prefrontal sources, reinforcing its downstream role in transforming prefrontal computations into affective or motivational outputs (Adolphs, 2010; Morrison & Salzman, 2010). This inhibitory prefrontal influence on the amygdala aligns with prior findings of top-down control in social contexts (Gangopadhyay et al., 2021; Putnam et al., 2023). Critically, dmPFC emerged as the sole origin of all outgoing dependency modulations, reinforcing its role as an upstream hub that generates predictions about others’ behavior and orchestrates downstream signaling within prefrontal and amygdala circuits (Apps et al., 2016; Báez-Mendoza & Williams, 2020; Hayashi et al., 2020; Iacoboni et al., 2005; Lockwood et al., 2016). The temporal cascade of modulations—early inputs to ACCg, followed by OFC, and ultimately the BLA—reveals a top–down, hierarchically structured information flow during social gaze. Further, our findings demonstrate the sensitivity of the information flow across the social brain circuits to social context (Jovanovic et al., 2022), such as those found for social decision-making (Dal Monte et al., 2020; Putnam et al., 2023) .

Additionally, our analyses point to dmPFC’s role as a central hub in the brain-behavior network, serving as a source of dependency modulations to all other nodes. This is consistent with the differentiated information flow patterns between dmPFC and BLA, in which dmPFC acts as the source of gaze content information to BLA during social gaze interaction (Qi et al., 2025). This observation also aligns with the well-documented role of dmPFC in Theory of Mind and social cognition, where it integrates contextual and emotional information to guide social decision-making (Báez-Mendoza et al, 2021; Báez-Mendoza & Williams, 2020; Hayashi et al., 2020; Iacoboni et al., 2005; Isoda, 2021; Martin et al., 2017, 2021). The strong modulatory influence of dmPFC on BLA and other prefrontal regions highlights its crucial role in shaping neural representations involved in social interactions.

Finally, the comparison of comprehensive and partial dependency modulation maps further validates the effectiveness of our super-session approach. The emergence of novel edges in partial maps, which could be explained by the comprehensive network, underscores the necessity of integrating data across multiple sessions to fully capture the complexity of brain-behavior interactions. This finding reinforces the idea that fragmented or single-session analyses may overlook critical dependencies that become evident only when a more complete dataset is reconstructed and viewed holistically.

Despite the utility of our approach, there are important limitations to consider. Due to recording constraints, we could only record two brain regions simultaneously during eye-tracking sessions. To construct a comprehensive network involving four brain areas and behavior, we developed a merging algorithm that aligns data across sessions into a super-session for DBN analysis. This algorithm was validated using simulated linear and nonlinear systems with known ground truth dependencies and further tested by swapping BLA activity across sessions. While effective, the F-score did not reach 1.0 (Fig. 2E), partly due to our use of a stringent scoring criterion and occasional identification of dependencies at incorrect time points—a known limitation of DBNs, especially in recurrent neural systems. Moreover, while partial graphs generated from raw data aligned well with the super-session network, some new edges did not conform to expected temporal patterns. For instance, when two edges emerged from a common node, the expected edge between downstream nodes after node removal was not always observed. These inconsistencies suggest that although the merging algorithm enables broader network reconstruction, temporal interpretations may be error-prone in recurrent systems. Future work incorporating multi-region simultaneous recordings or more sophisticated temporal models may help address these limitations.

Overall, our findings underscore the power of the brain-behavior network approach in revealing functional distinctions within highly interconnected neural systems and also emphasize the importance of interareal coordination in the prefrontal-amygdala networks in social cognition (Gangopadhyay et al., 2021). Previous work (Dal Monte et al., 2022) showed that regions such as the BLA, ACCg, OFC, and dmPFC all robustly encode social gaze variables—yet their individual contributions remained difficult to disentangle due to the extensive reciprocal connectivity and overlapping functional profiles of these prefrontal-amygdala areas (Kennedy & Adolphs, 2013; Joyce et al, 2025). In other words, when “everything is everywhere,” traditional analyses of single-region encoding provide limited insight into network-level specialization. By applying our DBN-based brain-behavior network framework, we were able to move beyond local representations and instead characterize how each region dynamically modulates or is modulated by others across time and context. This approach revealed nuanced functional differentiation—for example, the BLA’s temporally specific influence on prefrontal areas during different gaze roles, or the dmPFC’s hub-like role across conditions—that would be challenging to detect through isolated analysis of neural activity. More broadly, our results demonstrate that network-level modeling, especially in the context of social behavior, can yield critical insights into the distributed computations of brain systems where functional boundaries are blurred by dense interconnectivity. As such, the framework developed here offers a powerful tool for parsing the emergent properties of brain-behavior systems, particularly in domains like social cognition where interareal communication plays a central role.

## Acknowledgments

This research was supported by the National Institute of Mental Health (R21MH120672, SWCC, ASN, MPJ; R01MH110750, R01MH120081, and R01MH128190, SWCC), Simons Foundation Autism Research Initiative (SFARI 875855, SWCC, ASN, MPJ), Yale Orthwein Scholar Funds (ASN), and by the National Eye Institute core grant for vision research (P30 EY026878 to Yale University). We would like to thank Alec Sheffield for consultations on the computational modeling. We would like to thank the veterinary and husbandry staff at Yale for excellent animal care.

## Author contributions

ASN, SWCC & MPJ conceptualized the project. SF & ODM collected the data. FX analyzed the data. ASN supervised the project. FX, ASN, SWCC & MPJ wrote the manuscript.

## Supplementary Figure captions

**Supplementary Figure 1. Verification of “super-session” method by swapping BLA activity under different gaze conditions**

**(A)** Histograms of F-scores for the gaze pattern-matched BLA swapping group (cyan) and randomized BLA swapping control group (orange) for the condition in which the partner gazed at the actor. (**B**) Histograms of F scores for the gaze pattern-matched BLA swapping group (cyan) and randomized BLA swapping control group (orange) for the condition in which the partner did not gaze at the actor.

**Supplementary Figure 2. Probability distribution of matched behavior epoch durations**

The blue bars represent the histogram of matched behavior epoch durations, while the red curve depicts the exponential fit of their probability distribution.

## Methods

### Animals

Two adult male rhesus macaques (*Macaca mulatta*) were involved as recorded monkeys (Actor; monkeys L and K; aged 8 and 7 years, weighing 15.7 kg and 10 kg, respectively). A few animals served as partner monkeys (Partner). Over the course of the experiments using Actor-Partner pairings, Monkey L interacted with two adult male Partners and one adult female Partner (monkeys C, H, and E, all aged between 7 and 8 years, weighing 10.1kg, 11.1kg, and 10.7kg, respectively). Monkey K also interacted with two male Partners and one female Partner (monkeys L, H, and E). These resulted in six distinct macaque pairs for our behavioral and neuronal data (monkeys L-C, L-H, L-E, K-L, K-H, K-E). The actor and partner monkeys were unrelated and were housed in the same colony room with other macaques. In this study, all animals were kept on a 12-hr light/dark cycle with unrestricted access to food, but controlled access to fluid during testing. All procedures were approved by the Yale Institutional Animal Care and Use Committee and in compliance with the National Institutes of Health Guide for the Care and Use of Laboratory Animals. No animals were excluded from our analyses.

### Experimental setup

Each day, Actor and Partner were seated in primate chairs (Precision Engineering, Inc.) positioned 100 cm apart, with their heads 75 cm above the floor. Each monkey faced three monitors, with the central monitor placed 36 cm from their eyes. Two infrared eye-tracking cameras (EyeLink 1000, SR Research) continuously and simultaneously recorded the horizontal and vertical eye positions of both monkeys. Each monkey initially underwent a standard eye position calibration procedure, followed by an additional calibration to accurately map out the facial regions of each monkey (for more details see Dal Monte et al., 2022).

Each recording day included 10 social gaze interaction blocks involving a specific pair of monkeys. At the start of each session, the central monitors were remotely lowered to allow the monkeys to have an unobstructed view of each other. During the session, the monkeys were free to make spontaneous eye movements and interact with each other using gaze for five minutes. At the end of each 5-minute session, the central monitors were raised remotely, and the monkeys had no visual access to each other during a 3-minute inter-session break.

### Creating Balanced Behavior Variables

In our analysis, we examined instances of directed gaze between the two monkeys, specifically focusing on whether the Actor was gazing at the Partner. We defined this behavior as a binary variable: a value of “1” indicated that the Actor was gazing at the Partner, while a “0” denoted that the Actor was not directing gaze toward the Partner. However, in the raw dataset, this behavioral variable exhibited a substantial class imbalance, with the majority of time points corresponding to the Actor not gazing at the Partner (i.e., “0”). To address this imbalance and reduce potential bias in the modeling of gaze-related neural dynamics, we implemented a subsampling procedure to create a balanced distribution of “0” and “1” labels for Actor gaze. Specifically, we randomly sampled a subset of “0” time points to match the number of “1” time points, ensuring an equal representation of gaze and non-gaze events. This balanced dataset was then used in downstream dynamic Bayesian network (DBN) analyses to ensure that inferred dependencies were not skewed by class frequency.

### Merging algorithm

Our merging algorithm is based on matched patterns of gaze behavior. The eye positions of both monkeys were tracked every 1 ms. Gaze behavior was binned in 50 ms intervals using a Max-voting policy, such that the predominant behavior within each time bin was assigned to that bin. Using the generated behavior arrays, we developed an algorithm to identify the longest matched gaze patterns across different recording sessions involving various brain area combinations. Once the longest matched gaze patterns were identified, a super-session was created by merging the gaze variable and corresponding binned neural variables.

### Probability distribution and exponential fit of matched behavior epoch durations

After identifying the matched behavior epochs using the merging algorithm, their durations were plotted as histograms to illustrate the probability distribution (Fig. S2). An exponential fit was applied to characterize this distribution, yielding a time constant of 300.5 ms.

### Quantile scaling of neural data

Neural data was binned in 50ms intervals. To systematically analyze neural data across sessions with varying channel counts, we normalized the binned neural activity using quantile scaling (via MATLAB’s quantile() function) prior to input into the DBN pipeline. Sessions in which all quantiles were zero were excluded from analysis; this occurred in only one out of 25 sessions.

### Dynamic Bayesian Network pipeline

#### Fitting dynamic Bayesian network models

To infer dependencies among neural populations, a Dynamic Bayesian Network (DBN) framework was employed, utilizing a combination of the *pgmpy* Python package (https://github.com/pgmpy/pgmpy) and custom-developed Python scripts. Each binned and segmented data matrix underwent 100 bootstrap iterations. For each condition, a hill-climbing tabu search algorithm with a history window of three was applied to identify an optimal directed acyclic graph (DAG), initialized from a distinct random graph in each iteration. Model selection was guided by the Akaike Information Criterion (AIC). Variables at the most recent time slice (0 ms) were defined as effect variables, while all earlier time slices were considered potential causal variables. The resulting DAGs were unweighted, with edges representing binary (present/absent) dependencies. This procedure generated 100 unweighted DAGs per condition.

#### Estimation of weighted DAG

To estimate weighted DAGs, the 100 unweighted DAGs generated for each condition were subjected to an additional bootstrap procedure. Specifically, 100 bootstrap samples were drawn from the pool of unweighted DAGs, and edge-wise averages were computed. The weight of each edge, ranging from 0 to 1, reflected the proportion of DAGs in which that edge appeared, representing its consistency across iterations. This procedure resulted in 100 weighted DAGs per condition.

#### Testing significance of DAG edges

To evaluate the statistical significance of the identified edges, the 100 weighted DAGs per condition were compared against 100 control DAGs generated using an identical procedure but with temporally shuffled data (i.e., globally shuffled spiking activity). For each edge, a distribution of weights was constructed by aggregating values within the same condition. These empirical weight distributions were then compared to their shuffled counterparts using a one-sided Mann–Whitney U test. An edge was considered significant—and robust to time shuffling—if the weight distribution from the original (unshuffled) data was significantly greater than that from the shuffled data (p < 0.05).

### Simulations of linear and non-linear systems

In both the simulated linear and non-linear systems, X_1_ represents a random binary time series, while X_2_ is derived as a function of X_1_, and X_3_ is computed as a function of both X_1_ and X_2_. To test the performance of DBN in capturing the ground truth relationships between variables using different minimum matched pattern lengths, we generated super-sessions with varying minimum lengths. In these ground truth simulations, the X_1_ binary array was regarded as the anchor variable for pattern matching purposes. In the fully matched data, X_1_, X_2_, and X_3_ in the final super-session all obeyed the ground truth functional relationships. As controls, we created partially matched and completely unmatched data. In the partially matched data, either X_1_ and X_2_ or X_1_ and X_3_ in the final super-session followed the functional relationships; the remaining variable was randomly selected from its time series. In the completely unmatched data, X_1_, X_2_, and X_3_ in the final super-session do not have any functional relationships and were randomly selected from their respective time series. For each minimum match length, the simulation was run 100 times using independently generated X_1_ arrays to compute the mean and standard error of the mean.

### F score calculation

To evaluate how accurately the simulations matched the ground truth connectivity, an F-score was calculated for both linear and non-linear simulations, each fitted to their respective models. In this framework, edges were considered irrespective of their time lag. The F-score is defined by the given formula, where:

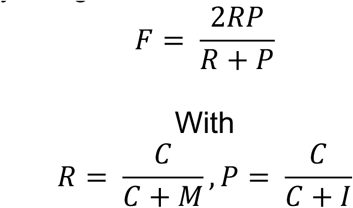

- C represents the number of connections correctly inferred by the simulation that are present in the ground truth,
- M represents the number of connections in the ground truth that were not inferred by the model,
- I represents the number of connections inferred by the model that are not present in the ground truth.

Recall (R) and Precision (P) are also incorporated into the calculation of the F-score.

### BLA swapping validation

Since BLA activity was recorded in every session, we swapped BLA activity between sessions with matched behavioral patterns. The original data served as the “ground truth”. After performing the BLA swap, we calculated the F-score between the swapped data and the ground truth. As a control, we also randomly sampled BLA activity from all sessions and calculated the F-score between this control group and the ground truth.

### Global shuffling

To perform the shuffling test, we developed an algorithm to randomize the raw spiking data. First, spike times were binned into 50 ms intervals to generate spike count arrays. These arrays were then temporally shuffled, and time points were randomly selected within each bin to reconstruct globally shuffled spike trains.

### Construction of prefrontal-amygdala dependency matrices

Data from the merging algorithm was fed into a DBN pipeline to construct a five-node dependency matrix during the “actor gazes at partner” condition. The five nodes consisted of the four prefrontal-amygdala brain regions and the partner gaze node. We then selected four brain-region-related nodes to create the four-node dependency matrix shown in Figure 5A. This entire process was replicated for the “partner gazes at actor” condition (four brain regions and the actor gaze node). The gaze nodes were included in the DBN fitting process to mitigate the detection of spurious dependencies between the neural nodes.

### Modulation index

The modulation index between two conditions in figures 4C and 5B is calculated as:

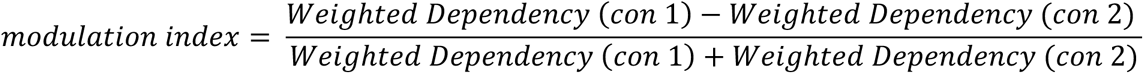

If the denominator is zero, then modulation index is set to zero.

