## Supplementary figures for "Causal Dynamics of Social Gaze in Primate Prefrontal-Amygdala Networks Revealed by Dynamic Bayesian Modeling"

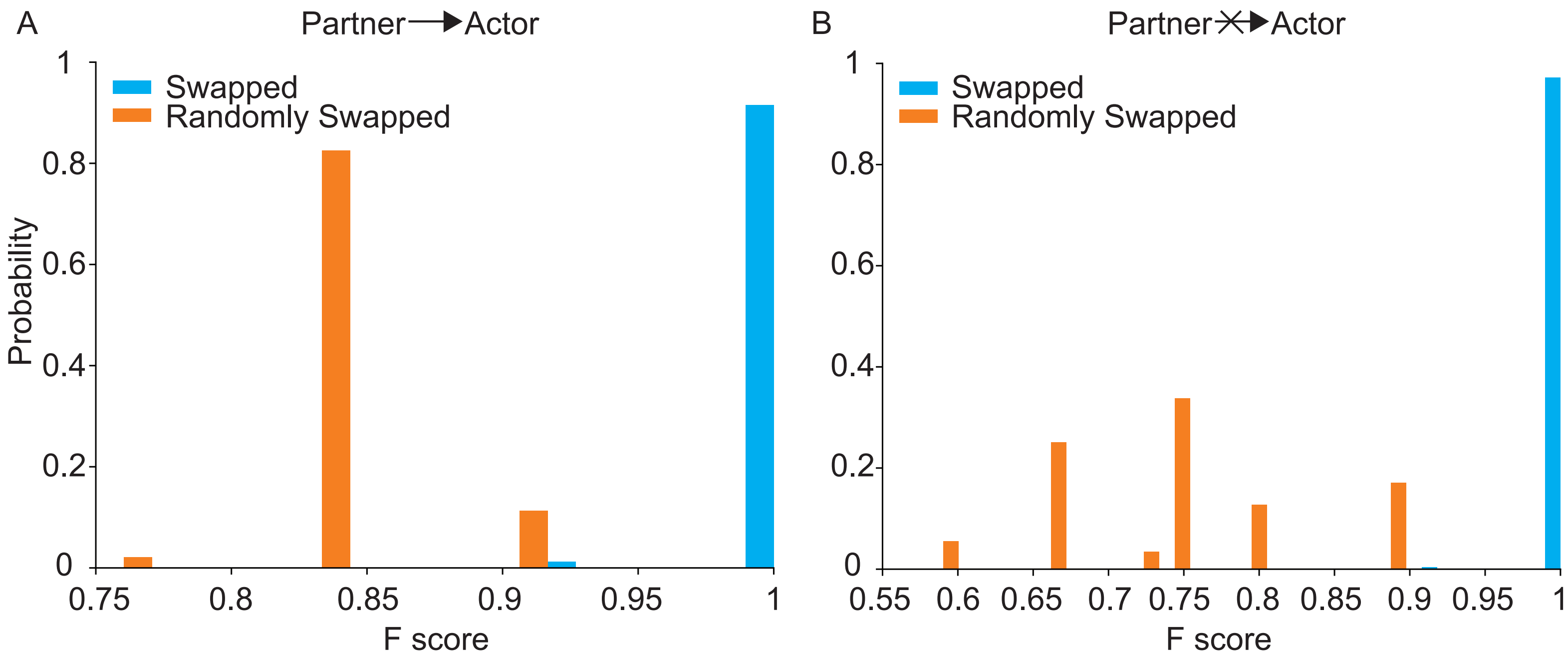


**Supplementary Figure 1. Verification of “super-session” method by swapping BLA activity under different gaze conditions**

(A) Histograms of F-scores for the gaze pattern-matched BLA swapping group (cyan) and randomized BLA swapping control group (orange) for the condition in which the partner gazed at the actor. (B) Histograms of F scores for the gaze pattern-matched BLA swapping group (cyan) and randomized BLA swapping control group (orange) for the condition in which the partner did not gaze at the actor.

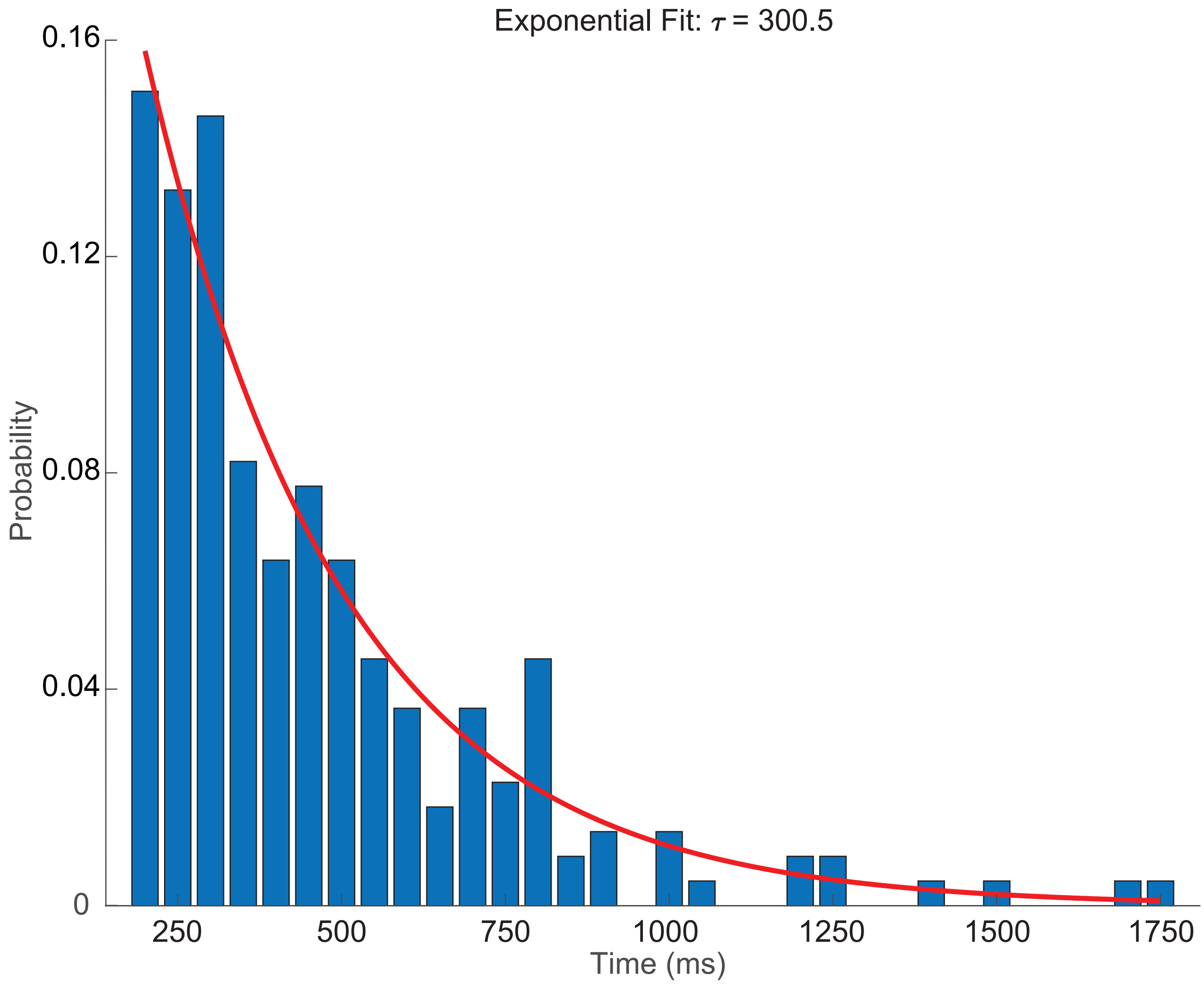


**Supplementary Figure 2. Probability distribution of matched behavior epoch durations**
The blue bars represent the histogram of matched behavior epoch durations, while the red curve depicts the exponential fit of their probability distribution.
